# A shared genetic basis for sexually antagonistic male and female adaptations in the toothed water strider

**DOI:** 10.1101/2024.03.27.586914

**Authors:** Claudia Pruvôt, David Armisen, Pascale Roux, Göran Arnqvist, Locke Rowe, Arild Husby, Abderrahman Khila

**Author notes:** Equal contribution (Claudia Pruvôt, David Armisen).

## Abstract

Sexual conflict can drive the divergence of male and female phenotypes and several cross-species comparative analyses have documented patterns of correlated evolution of sex-specific traits that promote the evolutionary interests of the sexes. However, male-female coevolution can be highly dynamic. Moreover, if male and female traits do not have an entirely distinct genetic basis, this can have profound effects on their coevolutionary dynamics. Here, we use water striders, a well-studied model system for sexually antagonistic coevolution, and ask whether sex-specific phenotypic adaptations covary across populations and whether they share a common developmental genetic basis. Using comparative analyses both at the population and species levels, we document an association between a derived male mate-grasping trait and a likely female anti-grasping counteradaptation in the toothed water strider *Gerris odontogaster*. Interestingly, in one population where males did not express their derived grasping trait, females had also regained the ancestral morphology. We then used experimental manipulation of gene expression, and show that these male and female traits are both linked to a common developmental genetic program containing Hox and sex determination genes, despite the fact that they are different structures on different segments. Our work thus suggests that the pleiotropic nature of developmental genetic programs can blur the distinction between inter- and intralocus genetic conflict.

## Introduction

Males and females share most of their genomes yet their evolutionary interests often diverge, resulting in evolutionary conflict (Arnqvist and Rowe, 2005; Bonduriansky and Chenoweth, 2009; Parker, 2006). We distinguish between intralocus (IASC) and interlocus (IRSC) sexual conflict (Schenkel et al., 2018). The former occurs when males and females have different optimal values for a shared phenotypic trait, which is manifested at a genetic level as different alleles in a given locus being favoured in males and females. Examples include height in humans (Stulp et al., 2012), age at maturity in salmons (Mobley et al., 2021) and body size in beetles (Berger et al., 2016; Kaufmann et al., 2021). A symptomatic pattern of IASC is a negative intersexual genetic correlation for fitness (Connallon and Matthews, 2019). In contrast, IRSC occurs when the optimal outcome of intersexual interactions differs in males and females. This can lead to the evolution of suites of sexually dimorphic traits encoded for by different loci in males and females that affect a given outcome, through a process known as sexually antagonistic coevolution (SAC) (Arnqvist and Rowe, 2005). Examples include the evolution of reproductive traits in seed beetles (Rönn et al., 2007) and molluscs (Koene and Schulenburg, 2005) and mate grasping traits in diving beetles (Bergsten and Miller, 2007; Bergsten et al., 2001) and water striders (Arnqvist and Rowe, 2002a, b).

Although SAC is thought to be a major driver of diversification and speciation (Arnqvist et al., 2000; Parker and Partridge, 1998; Rice et al., 2005), we actually know very little about the genetic basis of SAC (Pennell et al., 2016; Schenkel et al., 2018). This is unfortunate, because IRSC and IASC may not be entirely distinct if loci with effects on sex-specific traits involved in IRSC have pleiotropic effects with fitness consequences in the opposite sex (Geeta Arun et al., 2022; Mokkonen et al., 2016). If sexually dimorphic traits involved in SAC thus have, at least partly, a shared genetic basis, this will build a genetic correlation between male and female traits which significantly affects the predicted dynamics of SAC (Pennell et al., 2016).

In water striders (Gerridae), a monophyletic group of semi-aquatic Heteroptera, conflict over mating is widespread and is known to drive escalation of grasping traits in males and anti-grasping traits in females (Arnqvist, 1997; Arnqvist and Rowe, 2002a, b). In this SAC, males are favoured to mate repeatedly while multiple mating is superfluous and costly for females (Arnqvist and Rowe, 2005; Rowe, 1994). The outcome of male-female interactions is determined by pre-mating struggles, during which males strives to secure an anterior and a posterior grasp of the female, while female tries to dislodge the male (Arnqvist and Rowe, 2005). Sexually antagonistic traits that affect the outcome of struggles include modifications of the appendages or other body parts into claspers in the males, and the presence of spines, various integument projections and modifications of the tip of the abdomen in females that makes it more difficult for the male to grasp the female posteriorly (Arnqvist, 1989, 1997; Arnqvist and Rowe, 2002b; Crumière et al., 2019; Rowe et al., 2006). Phenotypic manipulation and behavioural studies have demonstrated the function of male and female sexually antagonistic traits during the pre-mating interactions (Arnqvist, 1989; Arnqvist and Rowe, 1995; Crumière et al., 2019; Han and Jablonski, 2009; Khila et al., 2012; Ronkainen et al., 2005), and a few studies have identified important genes required for shaping these traits during development (Crumiere and Khila, 2019; Khila et al., 2012).

Phylogenetic comparative analyses have revealed patterns of correlated evolution between the sexes under the influence of sexual conflict (Perry and Rowe, 2018). In water striders, Arnqvist and Rowe (Arnqvist and Rowe, 2002a, b) showed that the primary correlated changes are manifested in the shape of genital segments in both sexes, which determines posterior grip during struggles, and variation in sexually antagonistic traits correlates with variation in the economics of mating. Population level analyses also revealed that ecological factors, such as food resources, predation or population density, impact the patterns of antagonistic co-evolution of the sexes by altering the costs and benefits of mating (e.g., (Perry and Rowe, 2012; Rowe et al., 1994)). In theory, the dynamics of SAC in terms of escalation and de-escalation may thus be affected by both external and internal factors. First, environmental variation may act as driver of SAC and of phenotypic divergence between males and females (Perry and Rowe, 2012; Rowe et al., 2018). Second, the evolution of sexually antagonistic adaptations will affect their relative efficacy, and this in itself can generate escalation and de-escalation (Parker, 1979; Rowe et al., 2005). Third, the genetics of IRSC can itself contribute to the dynamics of SAC. In particular, between-sex pleiotropic constraints can act to generate cycles of escalation and de-escalation (Pennell et al., 2016).

It is therefore important to assess the genetic and developmental basis for sexually dimorphic traits involved in SAC. Here, we focus on a classic water strider species in this context – the toothed pondskater *Gerris odontogaster*. We first use comparative analyses at the micro- and macro-scale, to assess whether and how a well-known male grasping trait (consisting of two extensions of the seventh abdominal tergite) is matched by a female counter-adaptation. We include populations exhibiting significant variation, including the remarkable absence of the male grasping trait. Using experimental manipulation of gene expression, we then link co-varying male and female traits to the role of a developmental genetic program containing both Hox and sex determination genes.

## Results

### Correlated evolution of a male adaptation and a putative female counteradaptation

The *G. odontogaster* male abdominal processes (MAPs; Figure 1A) grasp the proctiger of the female during pre-mating struggles and thus play an important role in increasing mating rate by providing posterior grasp (Arnqvist, 1989). Females with a proctiger that is harder to clasp by MAPs should depress fitness costs associated with superfluous mating and, therefore, be favoured by selection. We here hypothesize that a concealed female proctiger may represent an adaptation in females to counter the effect of MAPs. This tenet is based on (i) comparative analyses in this clade showing negative correlated evolution between male clasping ability and the accessibility of the proctiger which affects mating rate (Arnqvist and Rowe 2002b), (ii) experimental manipulations showing that a less accessible tip of the abdomen makes it easier for females to dislodge males (Arnqvist & Rowe 1995, Ronkanen et al. 2005) and (iii) detailed studies of other semi-aquatic heteropterans showing that concealed genitalia represents a female counteradaptation (Han and Jablonski, 2009; Maroni et al., 2023). Populations of *G. odontogaster* differ considerably in mean length of MAPs, even on a small geographic scale, which is partly related to local population density and predation (Arnqvist, 1992b, 1994). Remarkably, males more or less completely lack MAPs in some populations. Such populations have been reported from Italy (Tamanini, 1979; Wagner, 1958), former Yugoslavia (Tamanini, 1979) and Mongolia (Lundblad, 1934), and have even been given the name *G. odontogaster brevispinis* (Lundblad, 1934) although such populations are thought to represent local types rather than a taxonomic unit (Andersen and Chen, 1993). To test our predictions, we phenotyped the females of three populations of *G. odontogaster*. First, we phenotyped a Swedish *G. odontogaster* population where males have fully formed and functional MAPs. In this population, the proctiger of the females is almost fully concealed under a dorsal plate consisting of a modified 8^th^ abdominal segment (Figure 1B). Second, we phenotyped *G. odontogaster brevispinis* from an Italian population where males have rudimentary MAPs (Figure 1C). In this case, we predicted that rudimentary MAPs should result in relaxed selection on the concealment of the females’ proctiger. Consistent with this prediction, the females of *G. odontogaster brevispinis* from Italy have a proctiger that is about four times more exposed compared to *G. odontogaster* from Sweden (Figure 1D, F). Third, we phenotyped *G. odontogaster* from France where males have MAPs that are of intermediate size between the Swedish and the Italian populations (Figure 1E). Interestingly, females also exhibited and intermediate state of proctiger concealment (Figure 1F). Finally, we phenotyped a population of *G. buenoi*, a closely related species to *G. odontogaster* and where males lack MAPs altogether. Again, the females of *G. buenoi* have an exposed proctiger (Figure 1F). A closer look on the females showed that the degree of proctiger exposure is primarily defined by the size of the 8^th^ abdominal segment, which consists of a dorsal plate on top of the proctiger. In *G. odontogaster* from Sweden, this plate is significantly larger and covers the proctiger, whereas this segment is narrower in *G. odontogaster brevispinis* from Italy and in other *Gerris* species allowing the proctiger to remain exposed (Figure 1B, D, F).

**Figure 1:**
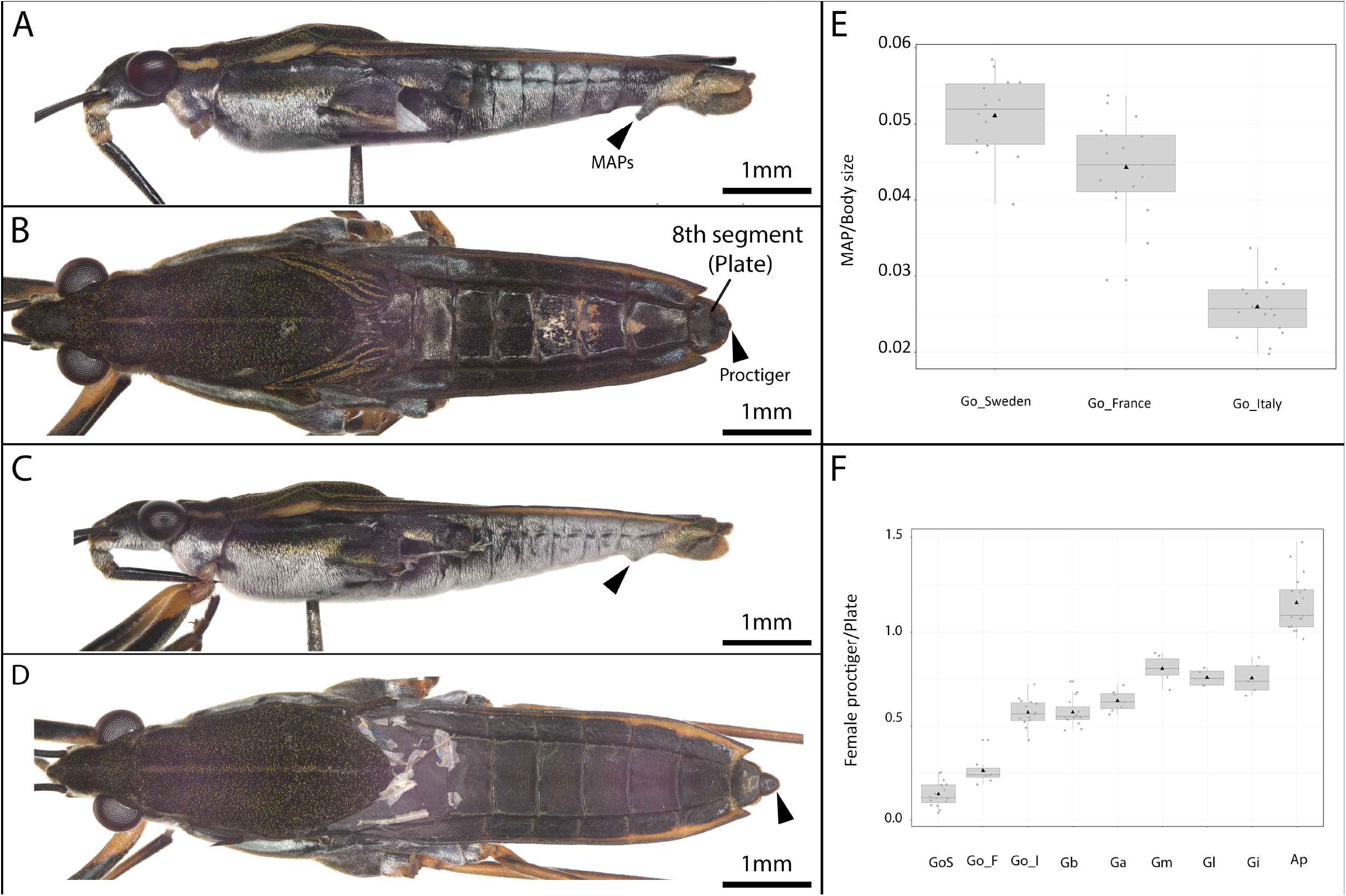
Phenotypic variation in the size of MAPs on the 7^th^ abdominal segment and the state of proctiger (eight abdominal segment) concealment in *G. odontogaster*. (A, B) male and female of a Swedish population. (C, D) Male and female of an Italian population. (E) Average size of MAPs in the Swedish, Italian and French populations. (F) Average proctiger concealment across *G. odontogaster* population and in a selection of *Gerris* spp.

Next, we wished to better place the above covariation between MAPs and the state of the proctiger across *G. odontogaster* populations in an evolutionary context. To assess the history of sexually antagonistic coevolution, we sampled four additional species of the genus *Gerris*, including *G. buenoi, G. lacustris, G. incognitus* and *G. marginatus* (Armisen et al., 2022). Females’ proctigers across all these species are fully exposed, consistent with the absence of MAPs in the males. A phylogenetic reconstruction of this sample based on the transcripts of 5 658 genes revealed that *G. odontogaster* from France and *G. odontogaster brevispinis* from Italy are sister populations and that *G. odontogaster* from Sweden is basal to both (Figure 2). In addition, this reconstruction placed *G. buenoi* as sister to all populations of *G. odontogaster* (Figure 2). This result confirms that the acquisition of MAPs is a derived state in *G. odontogaster*. Interestingly, this reconstruction also confirmed that the loss of MAPs in *G. odontogaster brevispinis* from the Italian population is a secondary event that followed their acquisition (Figure 2), providing evidence for a de-escalation of SAC. The evolutionary pattern of proctiger concealment is consistent with the state of the MAPs, suggesting that this concealment might be costly and that the ancestral state of proctiger exposure is regained by females when MAPs are lost in the males (Figure 2). To compensate for potential effects of phylogenetic dependencies, we tested for correlated evolution between the length of MAPs in males and proctiger concealment in females through the use of a phylogenetic generalized least squares (PGLS) regression. This model revealed significant negative correlated evolution (b = -0.079, SEb = 0.015, P = 0.003). These data further strengthen the suggestion that proctiger state in females is under selection by MAPs, generated by SAC.

**Figure 2:**
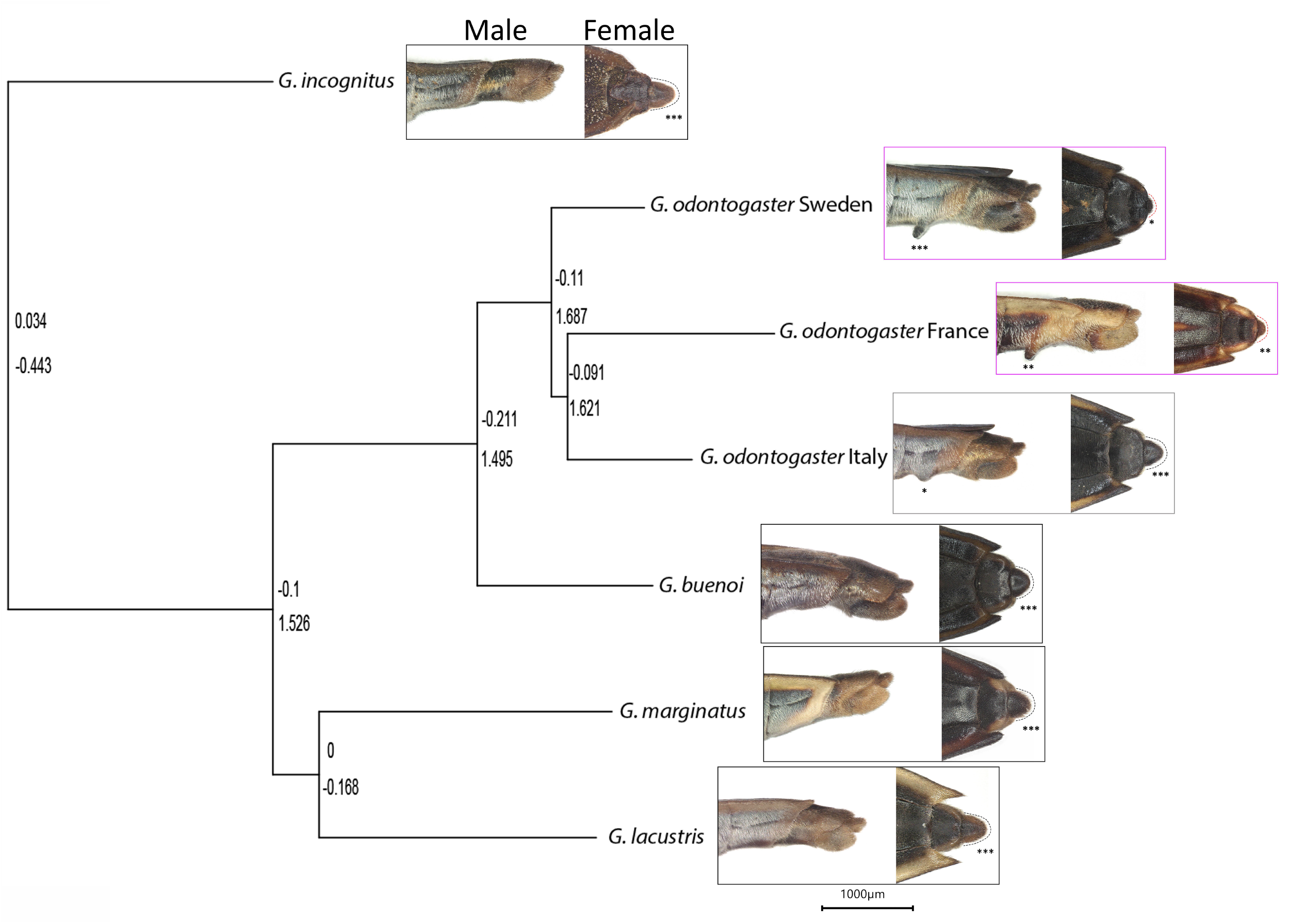
Phylogenetic relationships between three populations of European *G. odontogaster* and a sample of *Gerris* spp. Number of asterisks reflects the size of MAPs and the state of proctiger exposure.

### Cellular and developmental genetic basis of MAPs

Water striders belong to the Heteroptera suborder with post embryonic development consisting of five successive molts (Andersen, 1982). Phenotypic differentiation of males and females can only be observed in the genital segments at the fourth and fifth nymphal instars. The first signs of MAP specification appear at the fifth, and final, nymphal instar about 65 hours after molting from the fourth instar (Figure 3A, B). The developing MAPs can be observed through the cuticle of the 5^th^ instar nymph after treatment with alcohol (Figure 3A), or by peeling the cuticle of the nymph and uncovering the differentiating pre-adult integument (Figure 3B). MAPs development starts as a pair of cell populations organized into concentric circles that prefigure their position in the future adult (Figure 3B, C). The lumen of the developing MAPs is particularly rich in cytoskeletal material as revealed by actin and acetylated-alpha tubulin staining (Figure 3D). By the end of the fifth nymphal instar, MAPs extend and can be readily observed in the pre-adult after removing the nymphal integument (Figure 3C). These cellular and developmental dynamics suggest that the development of MAPs involves cytoskeleton remodelling and morphogenetic processes.

**Figure 3:**
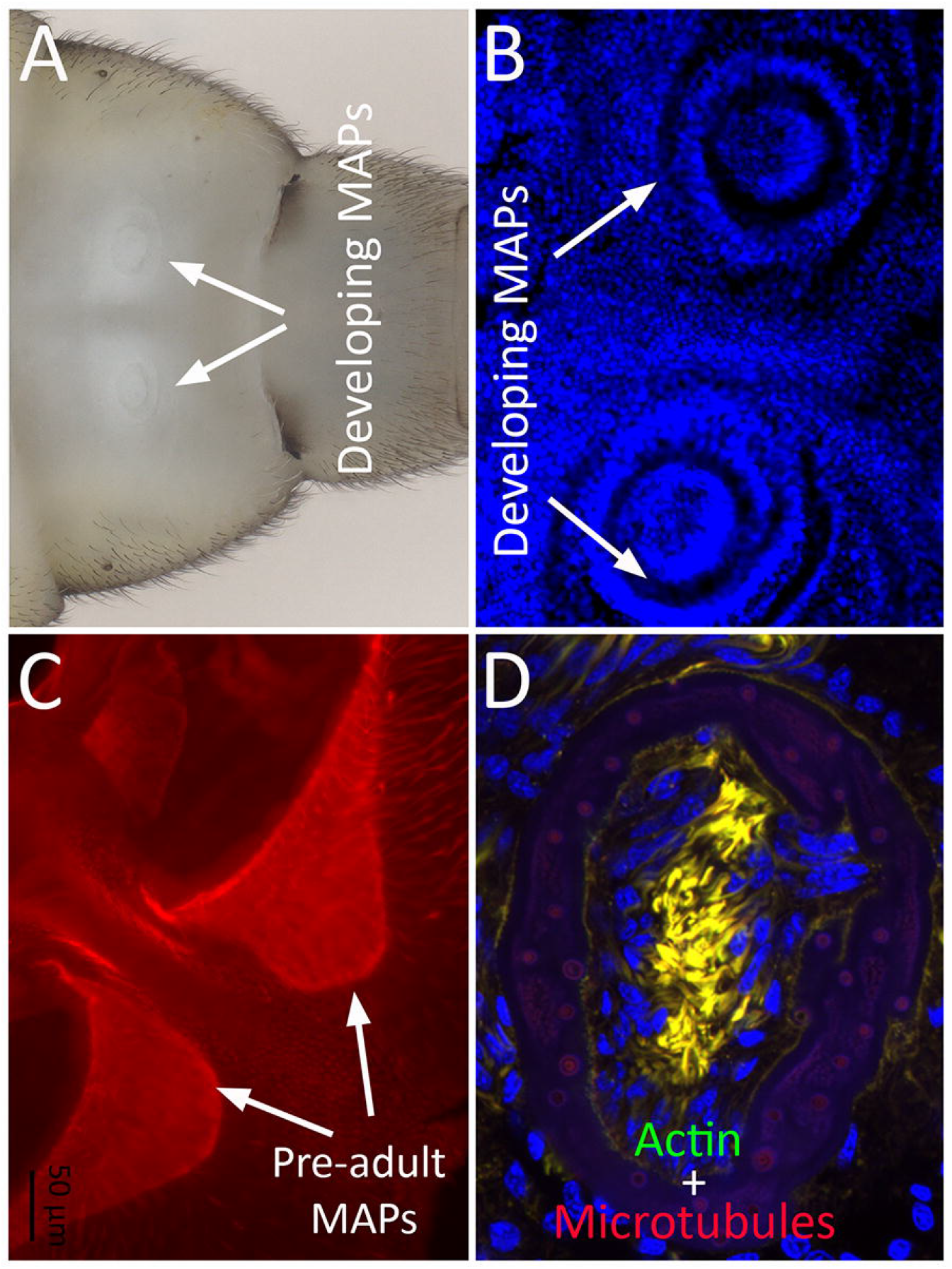
Development of MAPs. (A, B) The first signs of the development of MAPs appear at the middle of the fifth, and final, nymphal instar. (A) Two structures, prefiguring adult MAPs, can be visualized by imaging the nymphs in alcohol. (B) Epithelium of a pre-adult dissected out of the fifth instar nymph and stained with the nuclear marker DAPI. Developing MAPs consist of two sets of concentric cell populations which extend into full MAPs at the adult stage (C). (D) Actin (Green) and acytelated alpha tubulin (red) staining shows that the lumen of the developing MAPs is rich in cytoskeleton and microtubules. The yellow colour results from the overlap of actin (green) and tubulin (red).

### Hox and sex determination genes shape MAPs during development

The development of MAPs is genetically controlled (Arnqvist 1989), but the genes involved are not known. We hypothesised, based on the state of knowledge about the specification of abdominal segments in males and females, that the terminal Hox gene *Abd-B* and the sex determination gene *dsx* are involved in the development of MAPs. This is due to the position of abdominal processes on the male body and the fact that it is a sex-limited structure. Control males from the Swedish population, injected with buffer, were indistinguishable from wild-type males and MAPs were present and fully developed on the 7^th^ abdominal sternite (*n* = 7, Figure 4A-B). In wild-type and control males, the posterior part of this sternite had the shape of a trough that, we suggest, enables males to extend their genitalia in a more ventral angle in order to better reach females’ genitalia during per-mating struggles and mating (dashed outline in Figure 4B). Fifteen 4^th^ and early 5^th^ instar males injected with dsRNA targeted against *Abd-B* and 13 males injected with dsRNA targeted against *dsx* reached adulthood. Knockdown of both *Abd-B* and *dsx* resulted in the reduction or the complete loss of the MAPs (Figure 4C and E). In addition, knockdown of both genes resulted in the disruption of the trough-shaped posterior part of the 7^th^ abdominal sternite, which instead became more linear (dashed outline in Figure 4D and F). This overlap in the effect of *Abd-B* and *dsx* suggests that the two genes interact within a network that shapes male abdominal processes.

**Figure 4:**
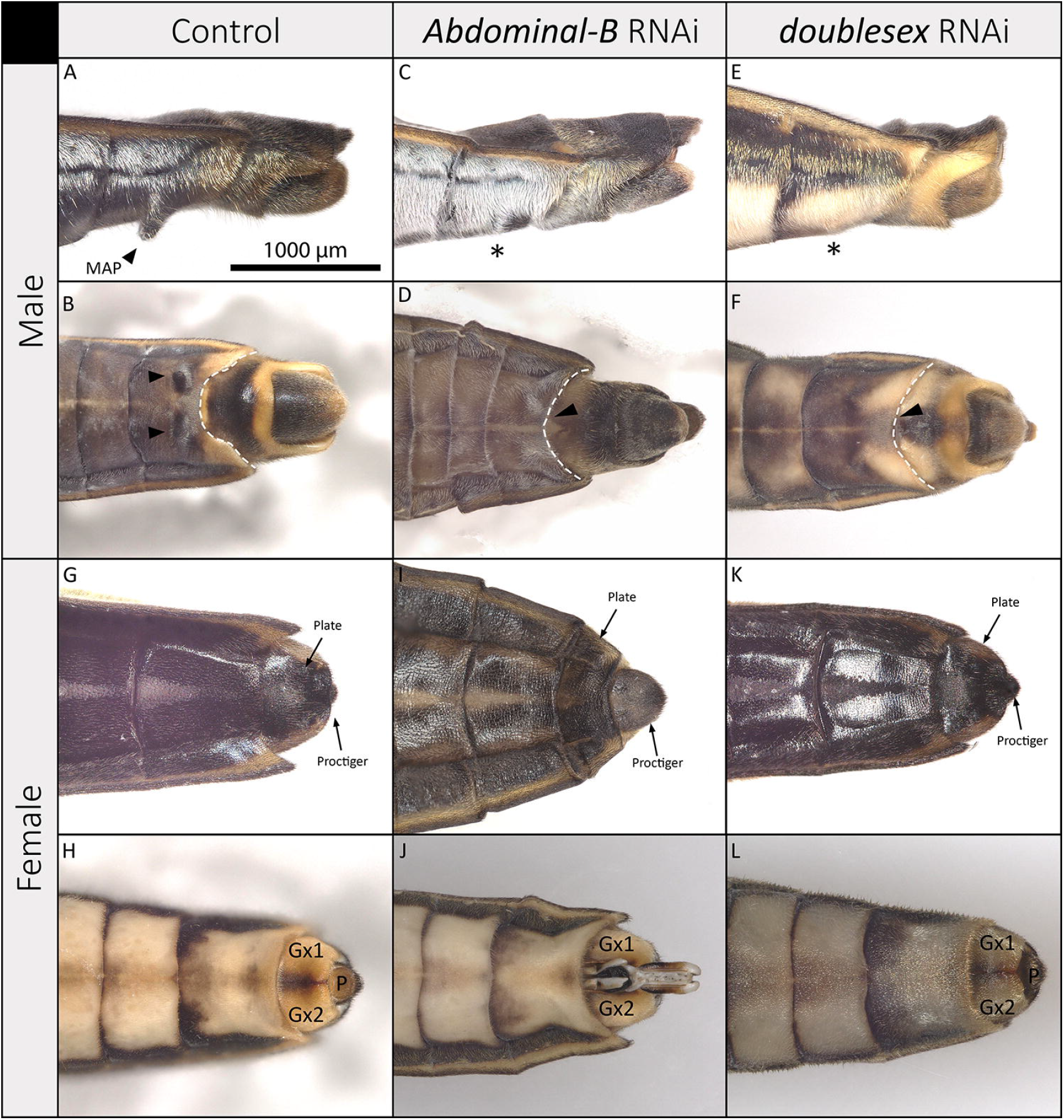
All images represent the Swedish population of *G. odontogaster*. Role of Abdominal-B and double-sex in the development of MAPs and proctiger concealment. (A) Lateral view of a control male showing fully extended MAPs. (B) Lateral view of a control male showing the trough shape of the seventh abdominal segment (dashed outline). (C, D) *AbdB* RNAi and (E, F) *dsx* RNAi treatments result in the complete or near complete loss of MAPs and the disruption of the trough shape of the segment (dashed outlines in D and F). (G, H) control females showing the state of proctiger concealment. (I) AbdB RNAi results in the narrowing of the plate that conceals the proctiger, leaving it exposed. (J) Other female genital structures, such as the gonocoxae, are also affected. (K and L) dsx RNAi does not seem to alter the concealment of the proctiger or the genital segments.

### *Abdominal-B*, but not *dsx*, is necessary for concealing female’s proctiger and shaping the genitalia

Control-injected 4^th^ and early 5^th^ instar females were indistinguishable from wild-type females (*n* = 7, figure 4G and H). 11 females reached the adult stage after the injection of *Abd-B* dsRNA. Interestingly, the 8^th^ abdominal segment that covers the proctiger became narrower leaving the female structure exposed (*n* = 11, Figure 4I). This suggests that *Abdominal-B* is part of a genetic mechanism that shapes female’s putative counter-adaptation through the concealment of the proctiger. The gonocoxae of wild-type females cover the ovipositor and the gonapophyses (Figure 4H). *Abd-B* knockdown resulted in a partial exposure of these structures (*n* = 7, figure 8C), and for two females they were completely exposed (Figure 4J). Adult females resulting from *dsx* knockdown had no clear morphological differences compared to controls (*n* = 16, Figure 4K and L). The absence of any effects of *dsx* knockdown in females suggests that this gene is not involved in the development of *G. odontogaster* female morphology during the final nymphal instar.

## Discussion

We have detected an association between a male sexually antagonistic trait (MAPs) and an apparent female counteradaptation (proctiger concealment) across several populations of *G. odontogaster*. These populations of *G. odontogaster* diverged recently from the North American species *G. buenoi*, which does not carry these traits. Moreover, our phylogenetic reconstruction showed that when MAPs are lost, the ancestral state of the proctiger (exposed) is regained. Importantly, we also provide experimental evidence for a link between these two coevolving traits: the Hox gene *AbdB* is involved in the development of both male and female sexually antagonistic traits, whereas the sex determination gene *dsx* are involved in the development of the male trait. Both genes are necessary for extending MAPs during development in males and their roles overlap significantly, whereas *AbdB* is required for the concealment of the proctiger in the female. We first discuss the coevolutionary pattern of these two traits, and then the developmental biology of these traits and its implications for SAC in this species.

Our analyses of the rapid coevolution of these sex-specific traits joins a small number of previous analyses of sexually antagonistic coevolution (Arnqvist and Rowe, 2002a, b; Bergsten and Miller, 2007; Bergsten et al., 2001; Koene and Schulenburg, 2005; Perry and Rowe, 2012). Previous phylogenetic comparative analyses in the genus *Gerris* have revealed correlated evolution between sex-specific and sexually antagonistic genital and pre-genital traits and showed that this is associated with the evolution of male-female interactions (Arnqvist and Rowe, 2002a, b). Our data mirror these findings at a microevolutionary scale, operating within the same species (see also (Perry and Rowe, 2012)), and further suggest that episodes of concerted escalation and de-escalation of sexual armaments can proceed rapidly in both sexes. This is the first clear example of de-escalation that we are aware of. We suggest that de-escalation is an expected outcome of sexually antagonistic coevolution. Parker (1979) showed that SAC may result in unresolvable evolutionary chases with escalation and de-escalation whenever male and female traits carry investment costs. In fact, models of sexually antagonistic coevolution that include natural selection on the antagonistic traits demonstrate that even where equilibria exist, changes in natural selection on both the traits, and the outcome of male-female interactions (i.e., mating) will shift those equilibria, causing both escalation and de-escalation (e.g., (Pennell et al., 2016; Rowe et al., 2005)).

Natural selection on these coevolving traits is likely occurring in *G. odontogaster*. In terms of MAPs, these are known to be costly to males in complicating the moult between the 5^th^ larval instar and adulthood (Arnqvist, 1994). This should increase the risk of cannibalism and predation during moulting, and the length of MAPs is indeed related to population density and predator presence across populations (Arnqvist, 1994). In terms of proctiger concealment in females, we suggest that this may also come at a cost. Since the proctiger primarily functions during egg deposition, trade-offs between efficacy as a resistance morphology versus egg deposition may render the proctiger sub-optimal in terms of attaching eggs efficiently on aquatic substrates.

Ecological impacts on the economics of sexually antagonistic interactions are well studied in water striders. Ecological factors, such as food abundance, predation, sex ratio and population density, are all known to affect the costs and benefits of sexually antagonistic traits in water striders (Arnqvist, 1994; Arnqvist and Rowe, 2005; Perry and Rowe, 2018; Rowe, 1994; Rowe et al., 1994). For example, food manipulation experiments in *Gerris* have shown that hungry females become more resistant to mating (Arnqvist, 1997; Rowe, 1992; Rowe et al., 1994) and high population density and male-biased sex ratios are both associated with relaxed sexual selection for long MAPs in *G. odontogaster* (Arnqvist, 1992a, b). Since *G. odontogaster* has a wide Palearctic distribution, the dynamic pattern of escalation and de-escalation of MAPs and proctiger concealment are consistent with variation in the ecological setting across its range.

Experimental manipulation of gene expression can provide information about the genetic programs shaping traits in both sexes, thus providing an important method to test whether correlated variation of male and female traits observed at the phenotypic level is linked to shared developmental genetic pathways. Depletion of two important developmental regulators, namely *AbdB* and *dsx*, resulted in the loss or severe alteration of male and female traits. This result is consistent with similar findings in *Drosophila melanogaster* (Kopp et al., 2000) and in *Photuris* fireflies (Stansbury and Moczek, 2014). Our work shows that MAPs in *G. odontogaster* males is developmentally at least partially controlled by a network including these two proteins. In flies, *AbdB* and *dsx* interact to establish sexual dimorphism in abdominal pigmentation (Burtis, 2002; Kopp et al., 2000). *AbdB* positively regulates the expression of *dsx* and it is a one-way interaction (Wang and Yoder, 2012). Here, *dsx* and another Hox protein (sex comb reduced) are known to regulate one another through a positive feedback regulatory loop (Tanaka et al., 2011). In our experiment, the effects of *AbdB* and *dsx* RNAi were very similar, as knockdown of either gene in males resulted in the loss of MAPs and the alteration of the trough-shaped 7^th^ abdominal tergite. This result indicates that the two genes interact and are part of a genetic network that shapes this male sexually antagonistic trait. The effects on two sex-specific male traits also show that these proteins have pleiotropic effects within males, which results in a genetic integration of MAPs and the shape of the 7^th^ abdominal tergite. Perhaps most importantly, *AbdB* RNAi also reduced the width of the dorsal plate leaving the proctiger exposed in females. This demonstrates that this protein is required for the development of both the male adaptation and its female counter-adaptation, a pleiotropic effect that should result in an inter-sexual genetic correlation between the traits, provided there is standing genetic variation in *AbdB*. A similar case of another Hox gene, *Sexombs-reduced*, involved in the development of male and female sexually antagonistic traits has been uncovered in the water strider *Rhagovelia antilleana* (Crumière & Khila, 2019).

Given that the male and female focal traits are different structures located on different segments, it would be easy to image them being underlain by independent loci. This is the common assumption of traits thought to be evolving independently and justifies the use of interlocus coevolutionary models in interpretation (Arnqvist and Rowe, 2005). However, we have shown that AbdB is necessary for the development of both of these traits. This result provides a rare empirical demonstration of a direct genetic link between IASC and IRSC (Mokkonen et al., 2016). If the observed evolutionary change in the two traits reflects either evolution of AbdB or its expression, then there is the potential for intra-locus conflict to play a role in their evolutionary dynamics. Theory has shown that a genetic correlation between male and female traits can act to de-stabilize SAC, especially when traits are costly (Härdling and Bergsten, 2006; Härdling and Karlsson, 2009). This is particularly true when the genetic correlation is caused by pleiotropy, blurring the distinction between IASC and IRSC (Pennell et al., 2016), when cyclic dynamics involving de-escalation in both sexes can occur. Alternatively, evolution of both of these traits may reflect evolution in the downstream targets of AbdB, and those targets may be independent of one another. A related example of this is the loss of abdominal pigmentation in *Drosophila santomea*, where both evolution of AbdB and its targets have played a role (Glassford et al., 2015; Liu et al., 2019). Our identification of a AbdB as a necessary component of the developmental network(s) underlying these rapidly coevolving traits enables more targeted study of the networks underlying their evolution and their degree of overlap.

## Materials and methods

### Study species

Water striders are hemimetabolous insects and undergo an incomplete metamorphosis and develop through five successive molts (Andersen, 1982). *Gerris odontogaster* individuals can be sexually differentiated at the 4^th^ nymphal instar (Supplementary Figure 1). The Swedish population of *G. odontogaster* was collected in 2019 in Uppsala, *G. odontogaster brevispinis* was collected in Castello-Molina di Fiemme (Italy) in 2022, and the population of *G. buenoi* was collected in a pond near Toronto (Canada) in 2010 and they have been bred in the laboratory since. Animals are fed daily with frozen crickets, and they are reared in water tanks in a room maintained at 28°C with a humidity between 40-50 % under an artificial photoperiod of 14 hours of daylight to mimic the reproduction season. Males and females are kept in the same water tanks to mate freely. Females lay eggs on Styrofoam floaters that are removed weekly and put into new water tanks to let the eggs hatch. A part of the French population was collected in a lake near Lyon in 2023 and the other part were collection specimen collected by Pierre Tillier in western France and by Olivier Durand in Maine-et-Loire (also western France). The selection of other *Gerris* species and the out-group species *Aquarius paludum* were collected in France and were stored in alcohol at -20°C.

### Phenotyping

Males and females from each species were photographed using a Keyence VHX 7000 microscope, then the animals were measured using software associated with the microscope. For males, the length of the abdominal processes and of the body were measured from a lateral view of the animal. For females, the visible part of the proctiger and the plate that recovers it were measured from a dorsal view.

### RNA interference against *Abd-B* and *dsx*

For this experiment, nymphs from 4^th^ and early 5^th^ instars of the Swedish population (largest MAPs) were injected with double-stranded RNA (dsRNA) of *Abdominal-B* and *doublesex*, separately, to induce degradation of transcripts corresponding to these genes (Agrawal et al., 2003). These nymphal stages were used insofar as individuals are sexually differentiated from the 4^th^ instar and it is before the beginning of the development of MAPs

The Cloning of *doublesex* and *Abdominal-B* was done by PCR based on sequences of *G. odontogaster* that were extracted from a full transcriptome database hosted at IGFL Lyon (https://gerromorpha.igfl.ens-lyon.fr/equipes/a.-khila-developmental-genomics-and-evolution/). Primers were designed based on these sequences using Primer-BLAST software (Ye et al., 2012). The cloning of AbdB and dsx fragments was carried out using standard PCR and cloning techniques.

### In vitro transcription of *dsx* dsRNA

Templates flanked by T7 promoters were synthesized by PCR using the primer sequences in (Lynch & Desplan, 2006). 1 µg of each T7 PCR template was used to synthesise double stranded RNA of *dsx* and *Abd-B* (Lynch & Desplan, 2006). 10 uL of 10X Transcription Buffer, 10 uL of 10 mM dNTPs, 1 uL of T7 RNA polymerase Plus (200 Units/uL), 1 uL of RNaseOUT ™, and a volume of nuclease-free water to obtain a final volume of 100 uL. The reaction was incubated at 37°C overnight: 2 uL of DNase (1 Unit/µL) were added to the reaction, which was incubated at 37°C for 10 minutes. Subsequently, the dsRNA preparation was purified using the RNeasy Mini Kit (Qiagen). Injection buffer (Rubin and Spradling, 1982) 10X was added (10 % of total volume). Finally, the dsRNA solution was filtered with Syringe Driven Filter Unit Cat, and was centrifuged at 5 000 rpm for 30 seconds.

### Injection of dsRNA of *dsx* and *Abd-B* into nymphs of *G. odontogaster*

Individuals from 4th instar to early 5^th^ instar of the Swedish population of *G. odontogaster* were used for this experiment due to the fact that the formation of MAPs begins 65 hours after the 5^th^ moult. Both sexes were injected to test whether or not the genes had a sex-specific function. Animals were anaesthetised with carbon dioxide for 15 to 20 seconds and then were injected between the 6^th^ and the 7^th^ abdominal segments with a 0.7 ug/uL dsRNA solution for *dsx* and 0.5 ug/uL for *Abd-B*. Each individual was injected with approximately 0.5 µL of this solution. Animals injected with the same volume of injection buffer 1X were used as negative controls to ensure that the trauma caused by injection does not induce a change of phenotype. Injections were delivered *via* quartz needles with filaments using a Zeiss Stereo Discovery V8 binocular magnifier. After injection, the animals were kept in separate water tanks and were fed on crickets twice a day until adulthood. After the injection of *doublesex*, males and females were kept separated in case the depletion of *dsx* transcripts had an effect on sex determination. When they became adults, individuals were sacrificed in ethanol, screened for phenotypes, and photographed using a Keyence VHX 7000 microscope. The experiment was terminated when at least 10 male adults per condition survived and had a modified phenotype.

### Fluorescence staining of MAPs

Mid-fifth instar male nymphs were dissected and the tergite of the 7^th^ abdominal segment with developing MAPS recovered. The tissue was fixed for 30 minutes in 4% paraformaldehyde, then washed five times in Phosphate Buffered Saline (PBS) containing 0.05% Tween-20 (PTW). Tissues were blocked for 1 hour in an antigen blocking solution containing 0.1% Bovine Serum Albumin, 1% Normal Goat Serum, 1X PBS and 0.05% Tween-20. The Tissues were then incubated with mouse acetylated alpha tubulin antibody overnight at 4 degrees Celsius. The tissues were washed five time, 10 minutes each, with PTW and incubated two hours with an anti-mouse antibody and Phalloidin (actin dye). Tissues were washed five times, 10 minutes each, with PTW and then five minutes in 1X PBS containing 30% Glycerol (in PBS). Tissues were incubated 10 minutes in 1X PBS, 50% Glycerol containing DAPI. Finally, the tissues were mounted on slides in 8% Glycerol. Images were captured using a Zeiss LSM 710 confocal microscope.

### Data analyses

Statistical analyses were conducted using R (v 4.2.2). The ratio of MAPs size/body size was compared between the males of the different *G. odontogaster* populations. The effect of population on this ratio was tested through a one-way analysis of variance (ANOVA). The normal distribution and homoscedasticity of the residuals were tested, respectively, through a Shapiro-Wilk test and a Bartlett test, and a graphical inspection of residuals was performed. When a significant difference was detected a Tukey post-hoc test was used to determine which populations differed. The same procedure was used to determine the impact of population and/or species on the ratio of proctiger size/plate size between females.

## Acknowledgements

We thank Pierre Tillier and Olivier Durand for sharing specimen of *G. odontogaster* (French populations). This work was supported by The Swedish Research Council (grants to GA and AK) and a CNRS MITI 80 prime PhD fellowship to CP.

